# Enteropathway: the metabolic pathway database for the human gut microbiota

**DOI:** 10.1101/2023.06.28.546710

**Authors:** Hirotsugu Shiroma, Youssef Darzi, Etsuko Terajima, Zenichi Nakagawa, Hirotaka Tsuchikura, Naoki Tsukuda, Yuki Moriya, Shujiro Okuda, Susumu Goto, Takuji Yamada

## Abstract

The human gut microbiota produces diverse, extensive metabolites which have the potential to affect host physiology. Despite significant efforts to identify metabolic pathways for producing these microbial metabolites, a comprehensive metabolic pathway database for the human gut microbiota is still lacking. Here, we present Enteropathway, a metabolic pathway database that integrates 3,121 compounds, 3,460 reactions, and 837 modules that were obtained from 835 manually curated scientific literature. Notably, 757 modules of these modules are new entries and cannot be found in any other databases. The database is accessible from a web application (https://enteropathway.org) that offers a metabolic diagram for graphical visualization of metabolic pathways, a customization interface, and an enrichment analysis feature for highlighting enriched modules on the metabolic diagram. Overall, Enteropathway is a comprehensive reference database and a tool for visual and statistical analysis in human gut microbiota studies and was designed to help researchers pinpoint new insights into the complex interplay between microbiota and host metabolism.

## Introduction

The human gut microbiota encodes a wide and diverse set of metabolic pathways for producing metabolites that can profoundly influence host physiology, health, and disease^1^. For instance, secondary bile acids such as Deoxycholate (DCA) and Isoallolithocholate (IsoalloLCA) are known to influence host metabolism^2^, cancer progression^3^, and the immune system^4,5^. These metabolites and their bacteria are potentially a great target for controlling the host phenotype in health and disease.

Together with investigating the bioactivity of metabolites produced by the human gut microbiota, there is an increasing number of experimental studies aimed at identifying the metabolic pathways and the genes involved in their production^6–11^. These studies allow us to extract the potential metabolic functionalities of metabolite production from the (meta)genomic sequence^12,13^ and enable the prediction of metabolite presence, leading to experimental investigations into their effects on host physiology.

Despite efforts to identify the metabolic pathways for producing microbial metabolites, a comprehensive metabolic database for the human gut microbiota is still lacking. Kyoto Encyclopedia of Genes and Genomes (KEGG) database, is one of the most widely used databases in metabolic studies, for many organisms, including bacteria^14^. However, the majority of the metabolic pathways in KEGG are based on those of humans and model organisms, and the specific metabolic pathways for human gut microbiota which are formed by multiple species, have yet to be covered^15^. To address this issue, biome-specific databases have been developed, like the human gut-specific metabolic modules^16^, and the gut-brain modules^17^ which are manually curated from scientific literature and were often used in recent human gut microbiota studies^18–21^. These databases encompass cellular enzymatic processes, which are defined as modules consisting of KEGG Orthology (KO) sets and the compounds involved in their production and degradation processes. However, they currently cover less than 160 modules, and they have yet to provide comprehensive coverage of microbial metabolic pathways such as secondary bile acids and trimethylamine (TMA) metabolism, and other important pathways. Furthermore, descriptions such as the relationship between microbial metabolites and/or metabolic pathways and the host information, including dietary patterns and diseases, are not provided. These descriptions can aid in generating hypotheses and revealing novel biological insights.

Here, we present Enteropathway, an integrated metabolic pathway database for the human gut microbiota, constructed through manual curation of the scientific literature. Its web application provides users with an interactive and customizable metabolic pathways diagram, as well as an integrated enrichment analysis module for highlighting enriched modules on the metabolic diagram. As a reference database and a visual and statistical analysis tool, Enteropathway is a valuable resource for human gut microbiota researchers on a quest for novel biological insights.

## Results

### Development of the Enteropathway database

Despite considerable efforts to map microbial metabolite production to the pathways of the human gut microbiota, a comprehensive metabolic database, specifically designed for the human gut microbiota is still lacking. Therefore, we developed Enteropathway, a metabolic pathway database that integrates 3,121 metabolic compounds, 3,460 chemical reactions, and 837 modules (a set of reaction processes) based on 835 manually curated scientific articles (**Figure 1, Table 1**). It covers microbial metabolic modules for producing/degrading diverse metabolites, including mono/oligo/polysaccharides, short-chain fatty acids, amino acids, pyrimidines, vitamins, bile acids, and other important metabolites.

**Table 1.** The database stats.

| Component | Overlap in KEGG | Overlap in gut-metabolic/brain modules | Only Enteropathway | Total |
| --- | --- | --- | --- | --- |
| Compound | 1759 | 0 | 1362 | 3121 |
| Reaction | 1395 | 0 | 2065 | 3460 |
| Module | 80 | 59 | 716 | 837 |

**Figure 1.**
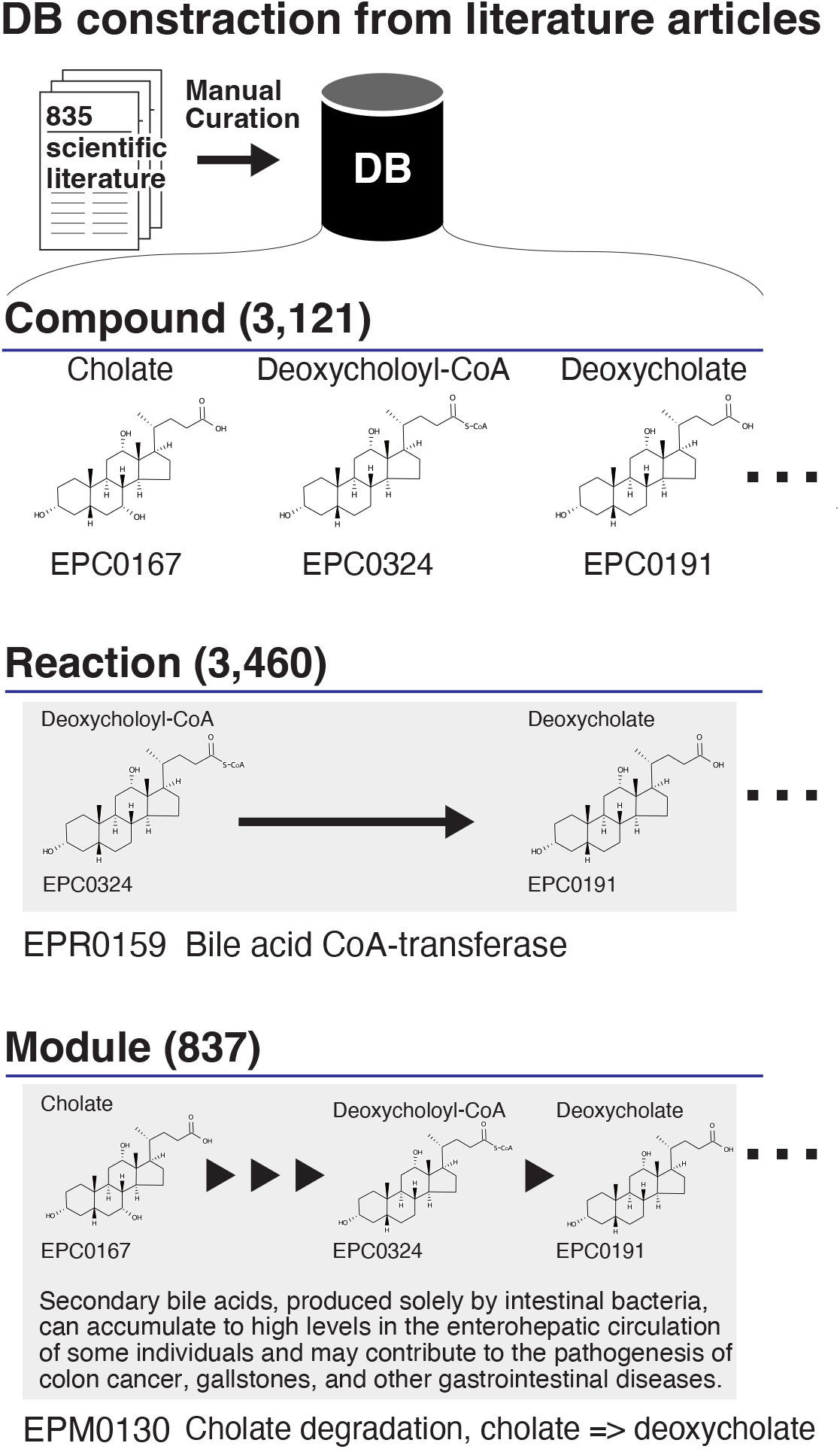
The overview of the database. The DB (database) was developed from literature articles by manual curation. This DB is composed of compounds, reactions, and modules. Each compound, reaction, and module is assigned an entry ID and a name. The reaction represents the chemical transformation for producing/degrading compounds. The arrow refers to the direction of the reaction. The module is generally defined as multiple reaction processes. The triangle represents one reaction in the module.

Subsequently, we investigated the relationship between these metabolic compounds and/or pathways and host-related information, such as dietary patterns and diseases. We then integrated this information into the description of 642 out of 837 Enteropathway modules. This enriched information can serve as a valuable resource for generating hypotheses and gaining novel biological insights.

Finally, we assigned unique Enteropathway identifiers to compounds, reactions, and modules, then manually linked them to widely used databases such as KEGG and UniProt by expert curation. This cross-referencing enables users to easily query Enteropathway with enzyme, reaction, or ortholog identifiers of commonly used databases in the field (**Table S1**).

### Development of the Enteropathway web application

Interactive metabolic pathway diagrams are powerful tools for understanding cellular metabolism^22,23^. When accessed through an interactive web application they can facilitate pathway exploration and visual analysis, enabling users to generate hypotheses and new biological insights. Therefore, we developed a web application (https://enteropathway.org), to provide users with a friendly way to interactively explore the Enteropathway database and its metabolic diagram.

Users can zoom in and out of the metabolic pathway with mouse gestures to comprehensively display the metabolic pathway at different levels, e.g. at the module level (**Figure 2a**). Additionally, the zoom buttons are also available for zooming in and zooming out as well as resetting to the initial zoom.

**Figure 2.**
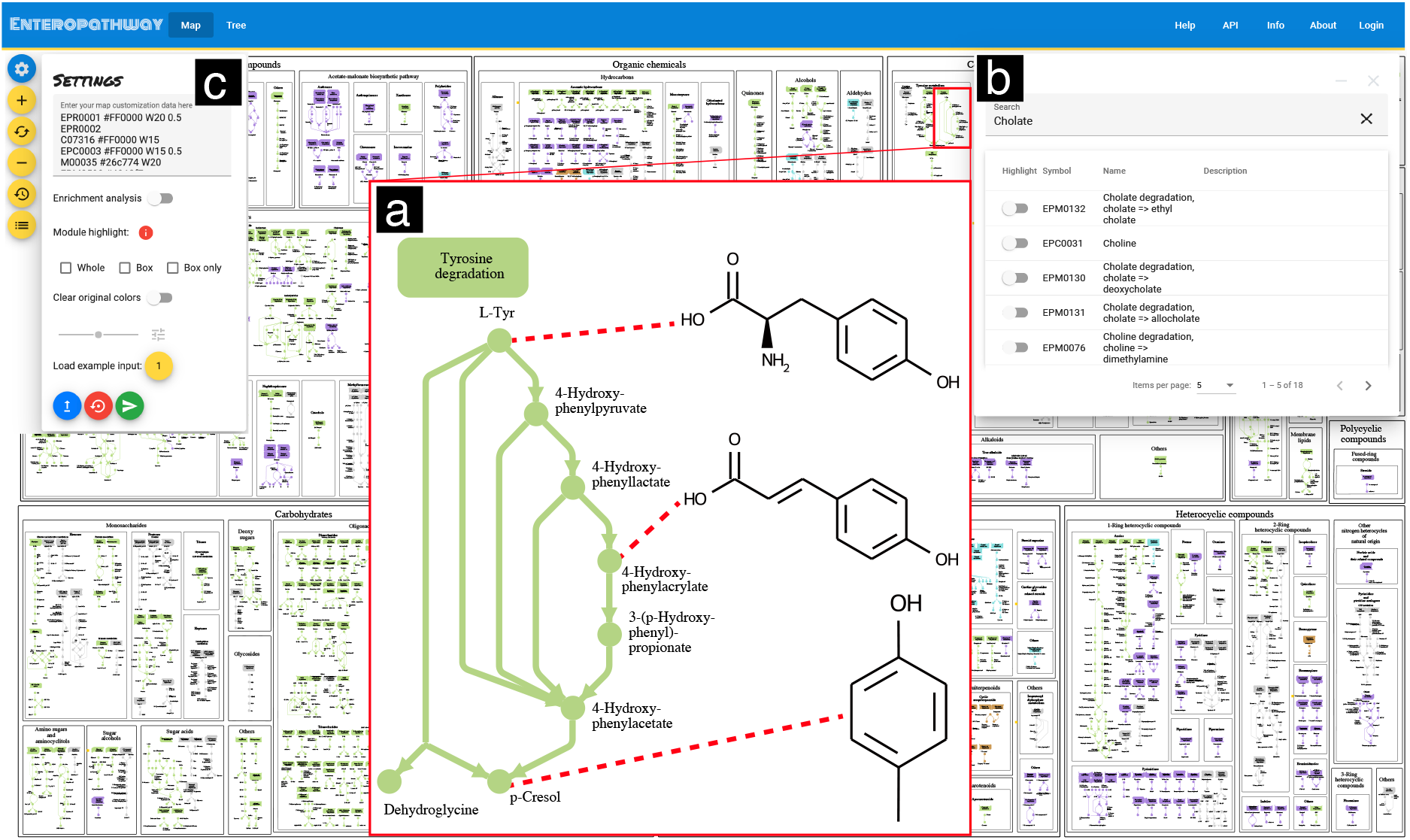
The user interface of the web application. The web application displays the metabolic pathway (**a**), search engine (**b**), and customization interface (**c**) in the following link https://enteropathway.org/#/diagram. (**a**) Each compound and reaction is displayed as a circle and arrow respectively. Each module is displayed as continuously concatenating circles and arrows, and its title is shown as the box. These are placed based on the module category in the metabolic pathway. (**b**) The search engine can explore the entry IDs by giving their entry names as keywords and then can highlight them on the metabolic pathway. (**c**) The customization interface allows users to characterize reactions, compounds, and modules on the metabolic pathway which are defined as their input data by clicking the submit button (green). The plus and minus buttons on the left side can zoom in or zoom out on the metabolic pathway respectively. The second yellow button on the left side can reset the initial zoom. The fourth yellow button on the left side can restore the original metabolic pathway.

The Enteropathway web application offers interactive features that allow users to conveniently access information related to the compound, reaction, and module. By clicking on the circle, arrow, or box on the metabolic pathway diagram, users can easily reveal the entry ID and access the corresponding entry page (**Figure S1, S2, S3**) which provides users with a list of manually curated scientific articles used as references, along with cross-references to other major biological databases. And to facilitate efficient exploration, Enteropathway incorporates a search engine that enables users to search and highlight their matched entities on the metabolic diagram, using entry names or IDs as keywords (**Figure 2b**). These functionalities should help users efficiently identify target metabolites or genes.

For large-scale exploration, a customization interface is provided for mapping analysis outcomes onto Enteropathway’s metabolic diagram (**Figure 2c**), using Entorpathway IDs or any IDs from the following list of supported annotations: KEGG (KO, Reaction, Module, Compound), Enzyme Commission number (EC number), EggNOG, MetaCyc Reaction, Rhea, Uniprot Accession, UniRef50, UniRef90, CAS ID. The interface accepts space-separated customization to target the color, size, and opacity (**Table S1**) of a metabolic pathway’s visual elements. Alternatively, the enrichment analysis module helps seamlessly highlight enriched reactions on the metabolic diagram (**Figure S4, see methods**). It accepts a list of KO or Enteropathway reaction IDs that will undergo a hypergeometric test to select and highlight statistically significant Enteropathway modules. The statistical test results can be shown and downloaded as a tab-separated format file. All diagram customizations can be exported as a PDF file, and user accounts are also available for saving and easily sharing customization results via a simple URL (**Figure S5**).

Finally, Enteropathway supports programmatic access through a REST API, allowing seamless integration with other bioinformatics tools. Detailed information can be found on the API page (https://enteropathway.org/#/api). Users can submit customization settings files via the API to obtain corresponding PDF results. Similarly, submitting lists of KO or Enteropathway reactions for enrichment analysis provides customized pathway diagrams in PDF format, along with statistical results in a tab-separated format file.

### Case study for analyzing metagenomic and metabolomic data sets by Enteropathway

The gut microbiota plays a role as a modulator of aging-related health by controlling immunosystems and resistance to pathogen infection^24–26^. A recent study showed that the secondary bile acids, especially Isoallolithocholate (IsoalloLCA) which was produced by Odoribacteraceae strains, were enriched in Centenarian (individuals aged 100 years and older) compared to Older (individuals aged 85-89 years) and Young (individuals aged 21-55 years)^*27*^. The IsoalloLCA may reduce the risk of pathogen infection by killing harmful gut bacteria such as *Clostridium difficile*. Here we use Enteropathway to further identify and visualize the Centenarian-specific genes and metabolites on its human gut-specific metabolic pathways diagram. To this end, we analyzed metagenomic and metabolomic data sets derived from the Centenarian cohort study^27^ and then explored them in Enteropwathway. Firstly, we downloaded these data sets and obtained KO, EC, and metabolome profiles (**Figure S6**). Then, we compared the abundance of KEGG Orthology (KO), Enzyme Commission (EC), and metabolites between Centenarian with Older and Young to identify Centenarian-specific reactions and metabolites. As a result, 3,884 KOs, 1,812 ECs, and 37 metabolites were significantly different in abundance in Centenarian (Q < 0.1, **Table S2**). These were converted to 1,291 reactions and 37 compounds and used to identify the Centenarian-specific module by enrichment analysis on the web application. This analysis showed that 67 modules were significantly enriched/depleted in Centenarian (Q < 0.1, **Table S2, Figure 3**). Among them, EPM0671, TMA biosynthesis, was enriched in Centenarian (Q=7.37×10^−5^). The TMA is the precursor of Trimethylamine N-oxide (TMAO), which is a well-known aging-related metabolite^28^.

**Figure 3.**
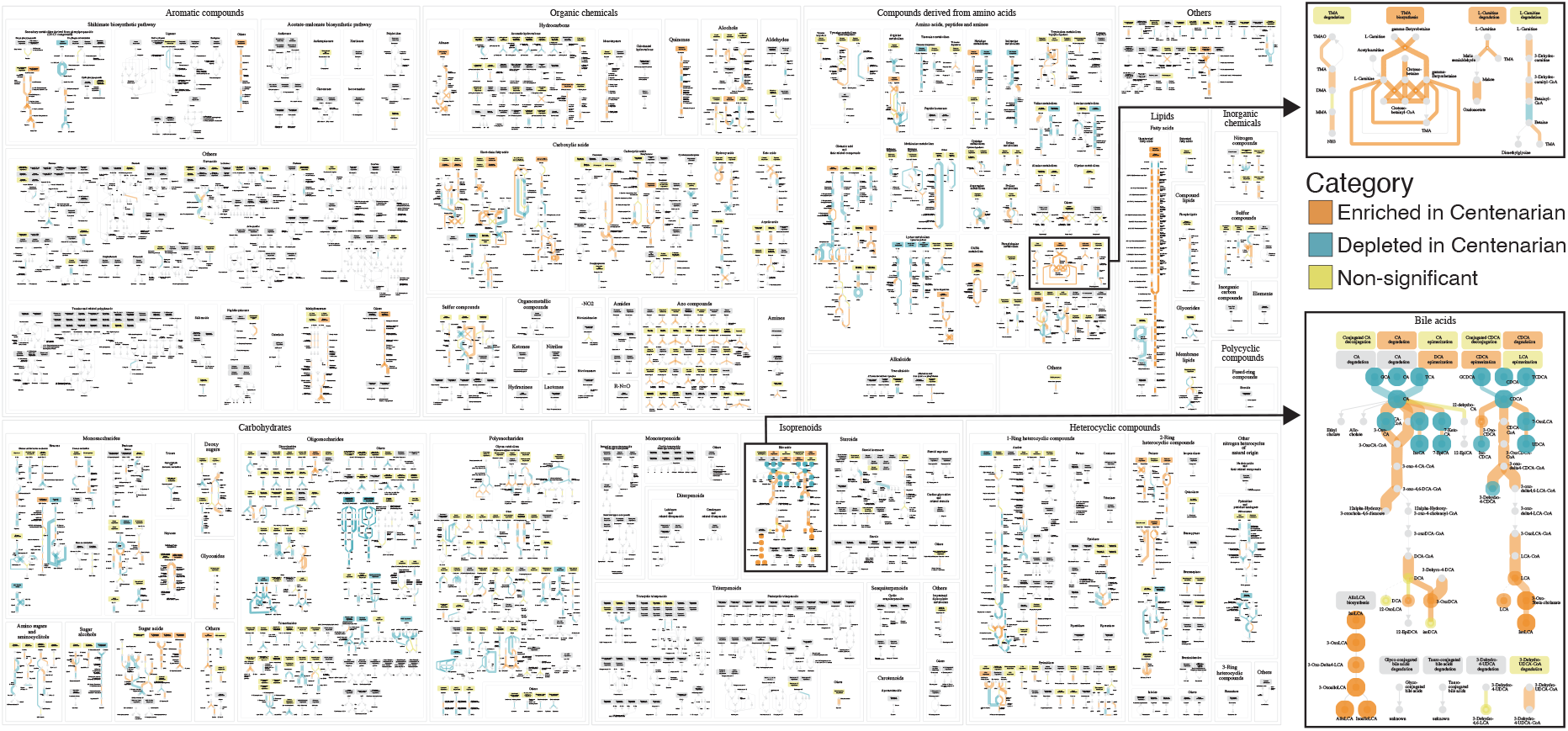
The metabolic pathway after customization of the metagenomic and metabolomic data. The metabolic pathway displays the enrichment of the Centenarian-specific compounds (circle), reactions (arrow), and modules (box). The shared link is https://enteropathway.org/#/diagram/-pNk6ieSu9H-EzmoppY72low9xcfmVXPjm3S4GK2xDvPggYXER1NoB4TKpQBYRHMlZZgcerzMmb2C10U7uEikeqqHUswmMoAvPl8d_E4RfEcHvCYtM_Dn7hxEWg9bqUERH3zw9XLSRM6eFJvCAPR_wYkvmXV40VBIh4f5HQBMu9BozVjOWtYUCvuFjWwlxoucSHuT78aOAHo3G_6RHaP96lLhpnCLKVSGCJ71286OvVer024N5tIWKqNG7f6zrpC. Centenarian-specific reactions, compounds, and modules are characterized by the Wilcoxon rank-sum test or the hypergeometric test. The color represents the significant differences (Q < 0.1; Orange: Enriched in Centenarian; Blue: Depleted in Centenarian; Yellow: Non-significant). The Q-value’s size reflects the reactions and compounds’ size and opacity.

We proceeded with customizing Centenarian-specific reactions, metabolites, and modules and specifically focused on the bile acids metabolism pathway for further analysis. By visually analyzing the customization result, we observed distinct enrichment patterns between the upstream and downstream sections of the pathway, providing valuable biological insights (**Figure 3**). In the upstream section of the pathway, Centenarian exhibited depletion of primary bile acids, specifically Glycocholate (GCA) and Taurocholate (TCA), along with their associated reactions. Conversely, in the downstream section, Centenarian showed enrichment of secondary bile acids, including Deoxycholate (DCA), Lithocholate (LCA), Isolithocholate (IsoLCA), Allolithocholate (AlloLCA), and Isoallolithocholate (IsoalloLCA), as well as their corresponding reactions. This observation suggests a reciprocal relationship between the depletion of primary bile acids in the upstream and the enrichment of secondary bile acids in the downstream. Further investigation of the final compound of the downstream in this pathway could provide valuable insights.

### Comparison with other databases and their visualization tools

KEGG and the gut-metabolic/brain modules^16,17^ have been used in many recent human gut microbiota studies^18–21^. To identify the set of novel pathways that are only covered by Enteropathway, we counted the number of unique Enteropathway modules that do not link to the KEGG database^14^ and the gut-metabolic/brain modules^16,17^. We found 1,362 out of 3,121 compounds, 2,065 out of 3,460 reactions, and 716 out of 837 modules that are unique to the Enteropathway database (**Table 1**).

Next, to evaluate the Enteropathway metabolic diagram, we customized Centanarian-specific reactions and compounds on the KEGG pathway (map00121, Secondary bile acid biosynthesis), then compared it to that of Enteropathway. We found that the upstream of bile acids metabolism was the same in KEGG pathways (**Figure S7**) and Enteropathway (**Figure 3**). However, the downstream was different as some of the compounds such as IsoLCA and AlloLCA are absent from the KEGG pathways, which suggests that Enteropathway can complement the KEGG pathway.

## Discussion

The human gut microbiota plays an important role in human health and disease. However, existing tools and metabolic databases for studying the human gut microbiota are limited. To address this gap, we have developed Enteropathway, an integrated metabolic database specifically designed for the human gut microbiota. It is based on an extensive set of manually curated scientific literature and contains several novel entries, especially metabolic modules that were never reported by commonly used databases such as KEGG^14^ and the gut-metabolic/brain modules^16,17^. This uniqueness positions Enteropathway as a valuable reference database for studies related to the human gut microbiota. The database is updated every three months and is accessible through a user-friendly web application.

In addition to pathway exploration, the web application provides users with mapping and enrichment analysis tools. Here we demonstrated these features by analyzing metagenomic and metabolomic data sets from a Centenarian cohort study. For instance, we could comprehensively capture the bile acids metabolism pathway and could find a difference in the enrichment between the upstream and the downstream within the pathway. This may reflect a bias for metabolic activity and presents the final compounds such as AlloLCA, and IsoalloLCA as potential targets for better understanding the difference between different age groups.

We further revealed the enrichment of potential metabolic functionality for producing TMA, the precursor of TMAO, in the Centenarian. This is consistent with another Japanese study^29^ that showed a high concentration of TMA in elderly subjects (individuals aged 60-85 years). Previous studies have shown TMAO is one of the cardiovascular disease and frailty markers^30,31^. Altogether, our result may suggest the high potential risk of these diseases in Centenarian.

The above shows how the Enteropwathy diagram is a powerful tool for finding biological insights at the metabolic level, and we hope it will lead other microbiome researchers to novel insights. Future enhancements include a statistical framework that will enable users to perform enrichment analysis directly from functional profiles. For the moment the user accounts system and the REST API provide different ways to analyze and easily share results from Enteropathway. Taken together, these features make our web application a user-friendly tool for pathway exploration, visual analysis, and statistical analysis.

However, it is important to consider certain limitations. Firstly, 52.4% of Enteropathway reactions are orphan enzymes. A computational approach to identify genes for orphan enzymes such as E-zyme2^34^, and their experimental validation are needed to characterize gene-based (meta)genomic data for human gut microbiota study. Secondly, in principle, modifications of microbiota-derived compounds by human enzymes such as hepatic enzymes are not covered in Enteropathway. Some microbial-host-derived compounds from these modifications have been reported as key risk factors for disease^35–37^. Therefore, focusing on these modifications is important to capture the impact of the gut microbiota on human health and disease.

In conclusion, Enteropathway is a comprehensive metabolic database for the human gut microbiota, curated from a wide range of scientific literature. It offers greater coverage of the metabolic pathway for the human gut microbiota compared to commonly used reference databases, positioning it as a valuable resource for studies in this field. Its web application provides an intuitive interface, allowing users to explore and customize reactions, compounds, and modules on a metabolic pathways diagram. Additionally, its enrichment analysis module enables the identification of potential metabolic functionalities and pathways involved in compound production or degradation. By analyzing publicly available metagenomic and metabolomic datasets with Enteropathway, we successfully uncovered enrichment patterns for aging-related compounds. This case study highlights how Enteropathway facilitates the discovery of biological insights through visual and statistical analyses.

## Methods

### Development of an integrated database for the human gut microbiota

The Enteropathway database for the human gut microbiota integrates compounds, reactions, and modules, derived from the manually curated scientific literature. Each reaction is assigned an Enteropathway reaction entry ID, along with a name and definition. Substrates and products resulting from the reaction are also assigned as an Enteropathway compound entry ID. Modules are defined as multiple reaction processes. Additionally, the database includes module descriptions that highlight the relationship between microbial metabolites, metabolic pathways, and host information, such as dietary patterns and diseases.

Each Enteropathway compound, reaction, and module entry ID have been manually linked to other databases by expert curation. Reaction IDs have been annotated with EC numbers^38^, Rhea^39^, KEGG Reaction, MetaCyc Reaction^40^, UniProt^41^, UniProt90, UniProt50, eggNOG^42^, and KO IDs. Similarly, compound entry IDs are linked to PubChem^43^, CAS (Chemical Abstracts Service), and KEGG Compound, while module IDs are linked to gut-metabolic/brain modules^16,17^ and KEGG Module IDs.

### Development of the web application

The Enteropwathy web application comprises a metabolic pathway diagram in SVG format, metabolic pathway browsers, and an enrichment analysis module. Each module in the diagram is color-coded according to its category, with green representing food, purple for drugs, brown for toxins, blue for secretions, and grey for other categories.

In the metabolic pathway browser, users can dynamically scale and customize the diagram using Enteropathway IDs or IDs from other databases which are matched and highlighted on the diagram. In case several IDs map to the same graphical element of the diagram, only the customizations of the last ID will be reflected on the diagram. Additionally, module customizations override reaction customizations within the diagram.

Several highlight options are available in the customization process. The color of the metabolic pathway can be changed to gray through the clear original colors button. The size of reactions and compounds can be interactively scaled by the slider. For modules, users can choose to highlight a module if at least one of its reactions is matched. They can additionally highlight the module title box or the module title box only. The enrichment analysis module identifies enriched or depleted modules based on a predefined library of reactions. Users provide a list of reactions as input, which are then linked to the corresponding modules. First, it counts the number of linked reactions in each module (referred to as A). Second, it counts the total number of reactions provided by the users (referred to as B). Third, it counts the number of reactions in each module (referred to as C). Fourth, it counts the number of reactions that are present in all modules (referred to as D). Finally, the platform compares the proportion of A to B with the proportion of C to D using the hypergeometric test from the ‘SciPy’ package in Python.

### Publicly available metagenomic and metabolomic data sets

We collected publicly available metagenomic and metabolomic data sets used in Sato et al ^27^ to analyze them with Enteropathway. In this study, 330 fecal samples were collected from 319 participants and classified into three groups according to age; (1) Centenarian (average age: 107 years old; 176 fecal samples derived from 160 participants), (2) Older (85-89 years old; 110 fecal samples derived from 112 participants), (3) Young (21-55 years old; 44 fecal samples derived from 47 participants). Samples derived from participants who underwent antibiotic treatment or had insufficient bacterial DNA yield have already been excluded.

For the metagenomic data sets, we downloaded 330 metagenomic samples from National Center for Biotechnology Information Sequence Read Archive (NCBI SRA, accession number: PRJNA675598). These samples were generated by Illumina NovaSeq 6000 with a 151 bp paired-end sequence (Number of paired-end reads per sample: 10 million).

The metabolomic data sets were obtained from the supplementary material of Sato et al^27^. In this study, 43 fecal bile acids were quantified from 297 metabolomic samples by LC-MS/MS (liquid chromatography-tandem mass spectrometry).

### Functional profiling

The functional profiles were generated by HUMAnN version 3.0.0^44^. In brief, metagenomic reads were mapped to the detected-species-specific pangenomes per sample, and unmapped reads were annotated by homology search against UniRef90 to derive UniRef90 gene family abundances in copies per million (CPM) units.

Subsequently, we obtained KO and EC profiles using the humann_regroup_table module from HUMAnN (--groups uniref90_level4ec and -- groups uniref90_ko) which maps UniRef90 gene families to KO or EC numbers, then sums the abundance of mapped families to obtain the abundance profiles. Finally, we normalized the profiles using the humann_renorm_table module with the parameter -- units relab to obtain relative abundances.

### Statistical analysis

A Wilcoxon rank-sum test was performed to characterize Centenarian-specific metabolites, KO, and EC by a comparison between Centenarian with Older and between Centenarian with Young. The Centenarian-specific modules were identified by a hypergeometric test using the Enrichment analysis module of the Enteropathway web application. To account for multiple testing, P-values were corrected using the Benjamini-Hochberg false-discovery rate, and Q-values were obtained. Statistical significance was defined as Q < 0.1.

### Pathway characterization

To characterize Centenarian-specific reactions, compounds, and modules on the metabolic pathway, we applied four methods to integrate statistics results derived from comparing KO and EC relative abundance and metabolites concentration between Centenarian and Older, and Centenarian and Young. Firstly, significant results derived from comparing either Centenarian and Older or Centenarian and Young were characterized as Centenarian-specific KO or EC or compounds. Secondly, Centenarian-specific KO and EC were converted to Enteropathway reactions.

Centenarian-specific compounds were also converted to Enteropathway compounds. Thirdly, Centenarian-specific modules were identified by enrichment analysis in the Enteropathway web application. Lastly, Enteropathway reactions, compounds, and modules were highlighted on the pathway by the web application with the option -- module_box_highlight box-only, --Clear_original_colors true.

Centenarian-specific Enteropathway reactions and compounds were converted to KEGG reactions and compounds respectively, then mapped on the map00121 in the KEGG pathway by KEGG mapper^23^ for comparing customization results between KEGG and Enteropathway.

## Acknowledgments

We are thankful to Dr. H. Mori for inspiring discussions. This work was supported by grants from the JST AIP Acceleration Research (JPMJCR19U3 to T.Y.), the Japan Society for the Promotion of Science (KAKENHI JP16H06279 (PAGS); JP25710016 (Grant-in-Aid for Young Scientists A) to T.Y.), the Japan Agency for Medical Research and Development (JP21ck0106546h0002 to T.Y.; JP21cm0106477 to T.Y; JP22ama221404 to T.Y.), the National Cancer Center Research and Development Fund (2020-A-7 to T.Y.).

## Author contributions

T.Y., S.G., S.O., Y.M., N.T., and H.S. contributed to the study concept and design. E.T., Z.N., and N.T. manually curated scientific literature to develop the database. Y.D., H.S., and H.T. developed the web application. H.S. performed bioinformatics analyses. H.S., T.Y., Y.D., T.N., E.T., G.S., and Z.N. wrote the manuscript. T.Y. supervised the study. All authors read and approved the final manuscript.

## Data availability

Nucleotide sequences of the Centenarian cohort from Sato et al are available in the NCBI SRA as PRJNA675598. The metadata and metabolomic profile for these samples are available in the Supplementary Table of Sato et al. The database as a tab-separated format file can be downloaded after login of the web application.

## Supplement figure legend

**Figure S1 The entry page of the compound**.

The entry page of the compound shows the Name, Description, Structure, References, and External references. This information is based on the database. The pink color represents that it can be accessed on different web pages by clicking. The link to the EPC0191 is following https://enteropathway.org/#/compound/EPC0191.

**Figure S2 The entry page of the reaction**.

The entry page of the reaction shows the Name, Description, Definition, Equation, References, and External references. This information is based on the database. The pink color represents that it can be accessed on different web pages by clicking. The link to the EPR0159 is following https://enteropathway.org/#/reaction/EPR0159.

**Figure S3 The entry page of the module**.

The entry page of the module shows the Name, Description, Reactions, Compounds, References, and External references. This information is based on the database. The pink color represents that it can be accessed on different web pages by clicking. The link to the EPM0130 is following https://enteropathway.org/#/module/EPM0130

**Figure S4 The user interface of the enrichment analysis**.

The web application displays the customization interface for the enrichment analysis (**a**) and analytic pane (**b**) in the following link https://enteropathway.org/#/diagram. (**a**) A set of reactions or KO can be used as the input type for enrichment analysis. The threshold of the P value can be scaled by the slider (default: 0.05). The module which is less than the threshold of the P value can be highlighted on the metabolic pathway by changing the color (default: red). The two options for highlighting the module are available (Box: highlight the module and title; Box only: highlights the module title box only). (**b**) The analytic pane displays the statistical results from the enrichment analysis based on the hypergeometric test. This result can be downloaded as a tab-separated format file by clicking the download button. The order of the module can be sorted in descending order of the P value or Q value by clicking the P value or FDR, respectively.

**Figure S5 The user interface for restoration and sharing**.

The web application displays the customization interface after login (**a**), the pane for restoration (**b**), and the pane for sharing (**c**) in the following link https://enteropathway.org/#/diagram. (**a**) After login, The download and the store function are available by clicking the second from the left button (blue) on the bottom and the first from the right button on the bottom (grey) respectively. The customization results can be stored nameable in the store step and can be restored later. The list for the restoration of the customization can be shown as (**b**) on the yellow button on the left side by clicking. (**b**) The restoration can be performed by clicking the purple button. The right button of the purple button can show the shared link as (**c**). The color of this button represents the shareable (red) or unsharable (yellow). (**c**) The yellow button can change from shareable to unshareable.

**Figure S6 The analysis pipeline of the centenarian cohort**.

The analysis pipeline shows the procedure of the customization result from the publicly available sequence reads, metadata sets, and the metabolome profile. The file and process are shown as the parallelogram and the square respectively. The color of the parallelogram represents the raw (black), intermediate (yellow), and final files (red).

**Figure S7 The customization results on the KEGG pathway**.

The metabolic pathway of the secondary bile acid biosynthesis derived from the KEGG pathway displays the enrichment of the Centenarian-specific compounds (square) and reactions (box) by the KEGG mapper. Centenarian-specific reactions and compounds are characterized by the Wilcoxon rank-sum test or the hypergeometric test. The color represents the significant differences (Q < 0.1; Orange: Enriched in Centenarian; Blue: Depleted in Centenarian; Yellow: Non-significant). The Q-value’s size reflects the reactions and compounds’ size and opacity.

## Supplement table legend

**Table S1 External link in the database**.

**Table S2 Statistical results**.

