## Supplementary material for "Enteropathway: the metabolic pathway database for the human gut microbiota": Figure S1

### EPC0191

Name: 3alpha,12alpha-Dihydroxy-5beta-cholanoic acid; Deoxycholic acid

Description

Structure

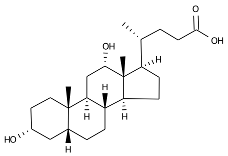


References

External references: CAS_ID:83-44-3
KEGG_COMPOUND:C04483
