## Supplementary material for "Enteropathway: the metabolic pathway database for the human gut microbiota": Figure S2

### EPR0159

| | |
|---|---|
| Name | bile acid CoA-transferase |
| Description | 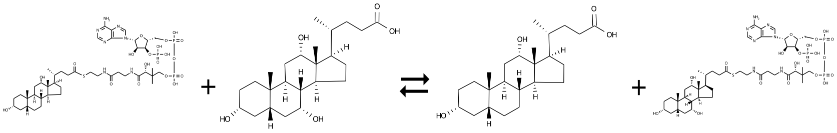 |
| Definition | Deoxycholoyl-CoA + Cholate <=> Deoxycholate + Choloyl-CoA |
| Equation | EPC0324 + EPC0167 <=> EPC0191 + EPC0318 |
| References | PMID:9990726<br>PMID:22021638<br>PMID:17764709 |
| External references | KEGG_KO:K15871<br>UniRef90:UniRef90_B4YST4<br>UniRef90:UniRef90_P19413<br>UniRef50:UniRef50_P19413<br>UniProt:P19413<br>UniProt:B4YST4<br>MetaCyc:MONOMER-18545<br>MetaCyc:BAIFEUBSP-MONOMER<br>Rhea:49439<br>eggNOG:COG1804<br>EC:2.8.3.25 |
