## Supplementary material for "Enteropathway: the metabolic pathway database for the human gut microbiota": Figure S3

### EPM0130

| | |
|---|---|
| Name | Cholate degradation, cholate => deoxycholate |
| Description | Secondary bile acids, produced solely by intestinal bacteria, can accumulate to high levels in the enterohepatic circulation of some individuals and may contribute to the pathogenesis of colon cancer, gallstones, and other gastrointestinal diseases. |
| Reactions | EPR0158 EPR1665 EPR0834 EPR0157 EPR0835 EPR0112 EPR2153 EPR0159 EPR3388 EPR3375 EPR1662 EPR0156 EPR2799 |
| Compounds | EPC0323 EPC0980 EPC0981 EPC0324 EPC2403 EPC1166 EPC0170 EPC0971 EPC1167 EPC1168 EPC0319 EPC0167 EPC0167 EPC0322 EPC0980 EPC0981 EPC0323 EPC2403 EPC0191 EPC0191 EPC0167 EPC2404 EPC0170 EPC0553 EPC0981 EPC0321 EPC0167 EPC1275 EPC0318 EPC0324 EPC0167 EPC0191 EPC0318 EPC0319 EPC2403 EPC0320 EPC0980 EPC0981 EPC0320 EPC0321 EPC2402 EPC0318 EPC2403 EPC0319 EPC0980 EPC0981 EPC0321 EPC0980 EPC0981 EPC0322 EPC2403 EPC1166 EPC0167 EPC0971 EPC1167 EPC1168 EPC0318 |
| References | PMID:16299351 |
| External references | |
