## Supplementary figures and images for "Enteropathway: the metabolic pathway database for the human gut microbiota"

### Figure S4

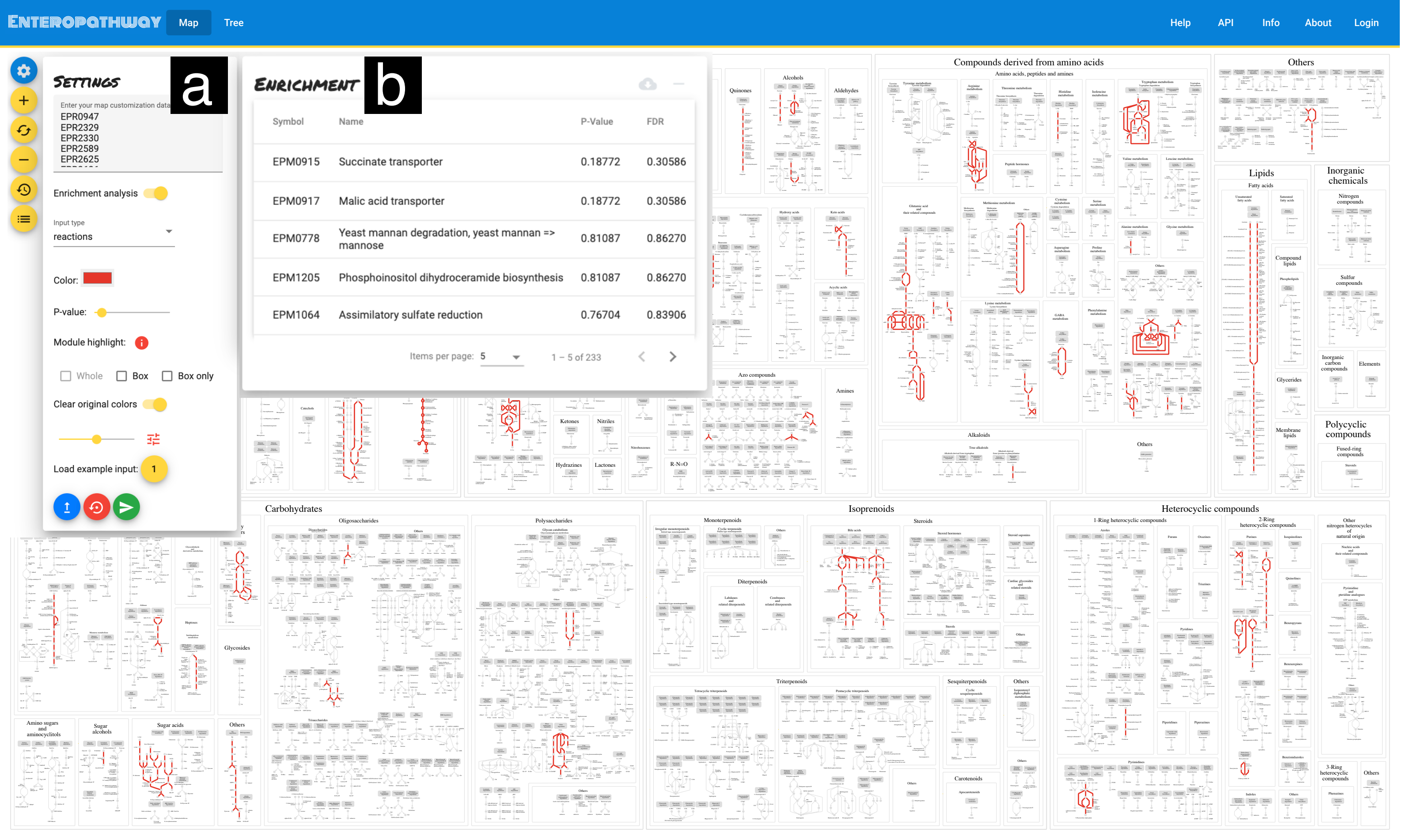

### Figure S5

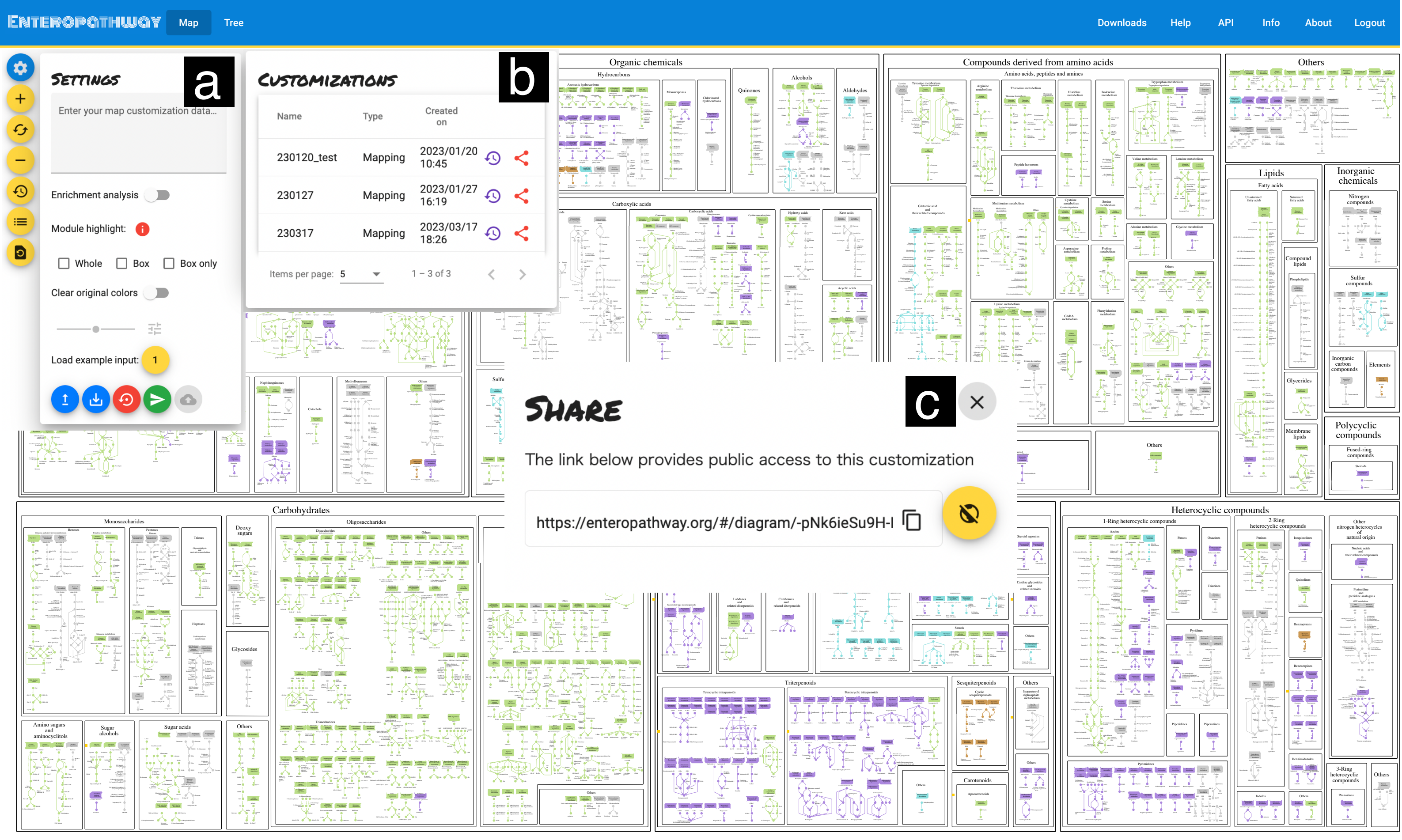

### Figure S6

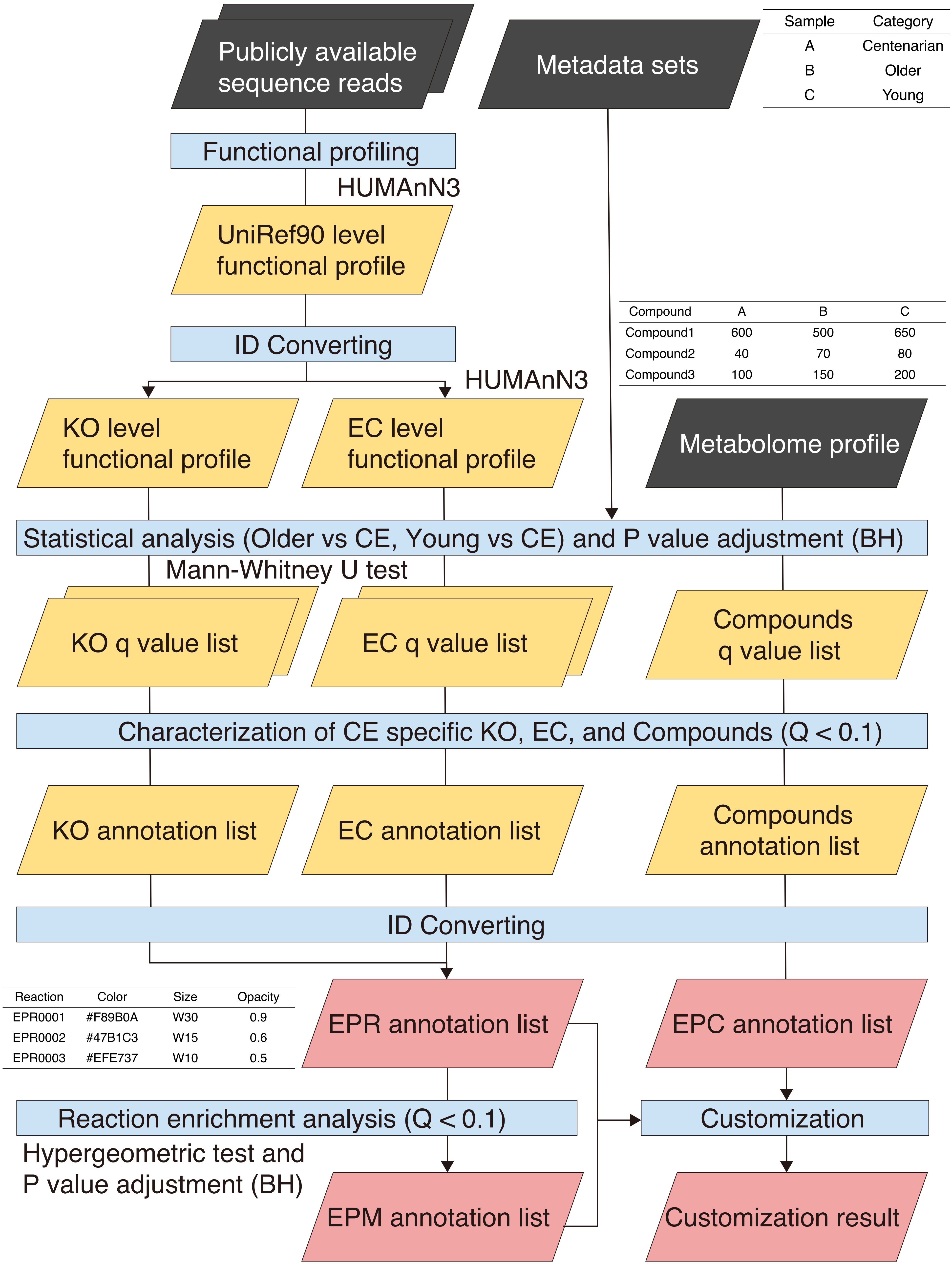

### Figure S7

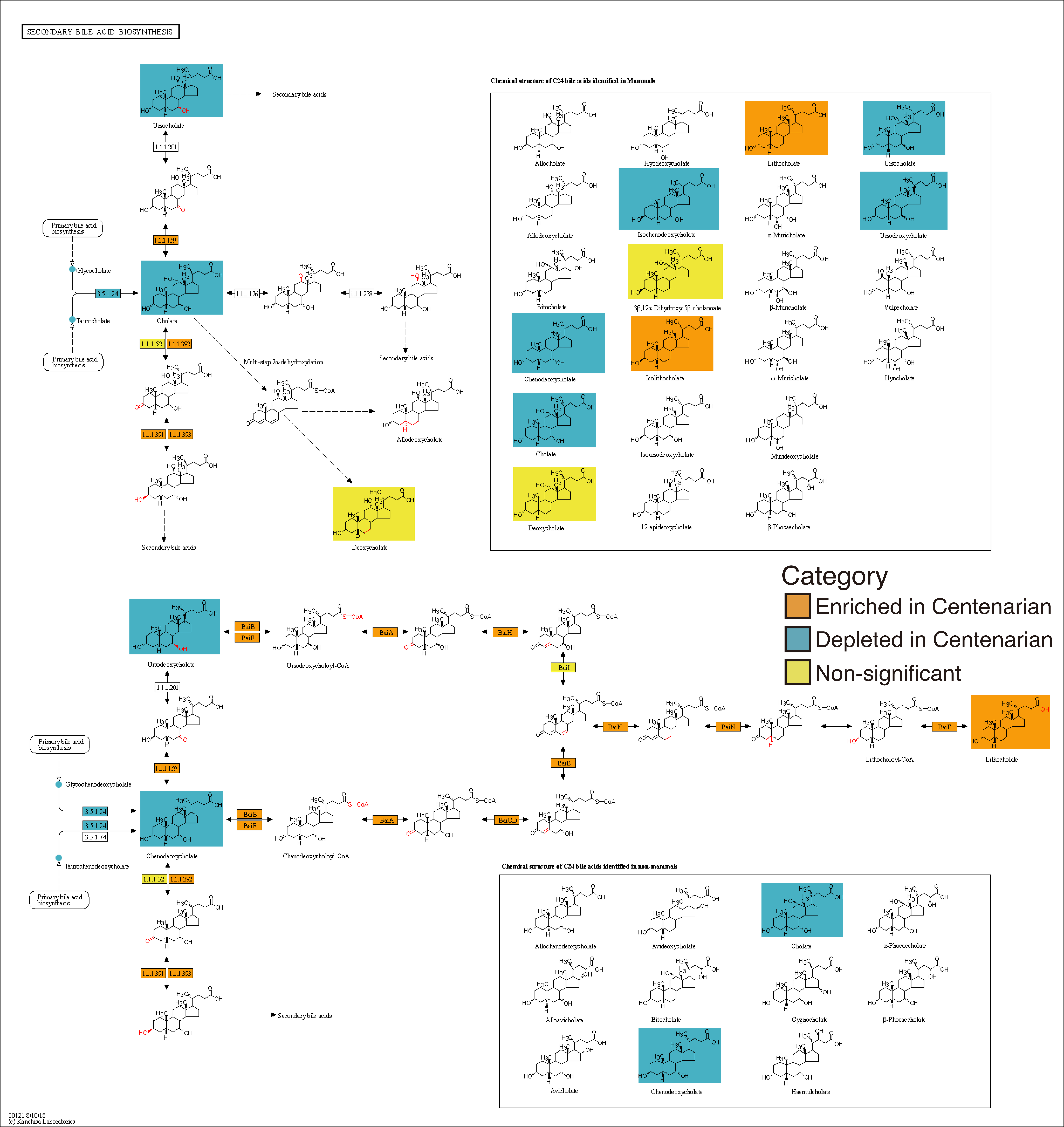
